# Multi-species numerical validation of an efficient algorithm for auditory brainstem response hearing threshold estimation

**DOI:** 10.1101/2024.02.19.580836

**Authors:** Erik Alan Petersen, Yi Shen

## Abstract

The Auditory Brainstem Response (ABR) can be used to evaluate hearing sensitivity of animals who are unable to respond to behavioral tasks. However, typical data collection methods are time consuming; decreasing the measurement time may save resources or allow researchers to spend more time on other tasks. Here, an adaptive algorithm is proposed for efficient estimation of ABR thresholds. The algorithm relies on the online update of the predicted hearing threshold from a Gaussian process model as ABR data are collected using iteratively optimized stimuli. To validate the algorithm, ABR threshold estimation is simulated by adaptively sub-sampling pre-collected ABR datasets for which the stimuli were systematically varied in frequency and level. The simulated experiment is performed on 5 datasets of Mouse (2 different datasets), Budgerigar, Gerbil, and Guinea Pig ABRs collected by different laboratories, with a total of 27 ears. The original datasets contain between 68 and 106 stimuli conditions, while the adaptive algorithm is run up to a total of 20 stimuli conditions. The adaptive algorithm ABR threshold estimate is compared against human rater estimates who view the full ABR dataset. The adaptive algorithm threshold matches the human estimates within 10 dB, averaged over frequency, for 19 out of 27 ears. The adaptive procedure is able to provide threshold estimates that are comparable to the human rater estimated thresholds while reducing the measurement time by a factor of 3 to 5. The standard deviation of threshold estimates from successive runs is smaller than the inter-human rater differences, indicating adequate test/retest reliability.

## 1. Introduction

The Auditory Brainstem Response (ABR) is an essential tool for determining hearing ability in populations that are unable to undergo behavioral assessment such as laboratory animals. The ABR is an evoked potential in response to an acoustic stimulus such as a click or tone-burst, which may be presented at different frequencies and sound pressure levels. A typical ABR measurement yields a time-domain waveform of the far-field electrical response from the brainstem to an acoustical stimulus. In general, as the stimulus presentation level is decreased, so does the amplitude of the ABR waveform. An ABR threshold may be defined as the lowest stimulus level for which a discernible ABR response is observed; if the stimuli are tone-bursts, a threshold contour may be mapped as a function of frequency.

Auditory brainstem responses are used for monitoring changes in hearing sensitivity *in vivo*, e.g., while testing the efficacy of a novel treatment for hearing loss, or a potential drug for ototoxicity (see, e.g., Zerche et al., 2023 and Stanford et al., 2021). In either case, ABRs allow the researchers to evaluate hearing sensitivity using physiological measurements.

One practical limitation of using ABRs to evaluate hearing sensitivity is the considerable duration of a typical measurement, which may last 30-45 or minutes or longer per ear (Schrode et al., 2022). The time required for ABR measurements are limited by two main factors: (1) the number of times that a stimulus must be repeated to obtain a synchronously averaged ABR waveform, and (2) number of stimulus conditions that are needed for the estimation of ABR thresholds across frequencies. To address this, a number of methods have been developed to reduce the total time of ABR measurements. One approach is to use *F* -ratio type quantities such as the *singlepoint variance*, which calculates the variance of a single data point across repeated measurements during synchronous averaging to determine the likelihood that a signal is present in noise (Elberling and Don, 1984). Another approach is to devise methods to increase the rate of stimuli presentation without causing neural adaptation, e.g., through deconvolution of responses at high presentation rates (Delgado and Ozdamar, 2004, Kaf et al., 2017 Mar/Apr), interleaving frequencies (Buran et al., 2020) or presenting stimuli to both ears simultaneously (Polonenko and Maddox, 2022 Mar/Apr). These methods increase measurement efficiency on the order of 2-3 times compared with traditional procedures.

In laboratories studying the auditory system using animal models, ABRs are often measured following a long list of pre-determined stimulus frequencies and levels. However, because the researchers may have no *a priori* knowledge of where the threshold is, a large number of these stimuli conditions will be well above, or well below, the threshold, and therefore are not necessary for threshold detection. Time-saving could be achieved by using only the stimuli conditions that provide the most information gain for threshold estimation, rather than following a longer pre-determined list of stimuli conditions.

Bayesian adaptive algorithms present one approach for efficiently estimating characteristics of a system, such as ABR thresholds, through optimized stimulus selection. Stimulus optimization is accomplished by fitting a model to the data collected thus far; the model can then generate predictions for the outcome of a future trial, which can be used to inform the sampling strategy for maximum information gain. The model can be parametric, with explicit formulation of the underlying psychophysical function, or non-parametric, for which limited assumptions are imposed on the formulation of the underlying psychophysical function.

Bayesian adaptive stimulus selection has been used widely in electrophysiology. It has been used to optimize the stimuli of implanted devices, e.g., for spinal chord stimulation for some patients with spinal chord injuries (Zhao et al., 2021), and for deep brain stimulation devices intended to treat motor symptoms of Parkinson’s disease (Grado et al., 2018). In both of these cases, the utility of a Bayesian approach is to maximize a measurable positive response to the treatment, such as restored mobility. Bayesian adaptive stimulus selection has also been used to characterize neural responses, such as the input and out curves for transcranial magnetic stimulation (S. M. M. Alavi et al., 2019). For these studies on humans subjects, the utility of a Bayesian adaptive stimulus selection scheme is often for increased efficiency to reduce overall testing time. See Pillow and Park, 2016 for a review of Bayesian adaptive procedures developed to measure neural tuning or response characteristics.

Adaptive sampling strategies that update stimulus selection schemes has been used throughout psychophysics, in particular, for vision (see e.g., Watson, 2017, Song et al., 2018, Lesmes et al., 2006, Lesmes et al., 2010, Dorr et al., 2017, and Marticorena et al., 2024). Bayesian adaptive sampling has also been successfully applied in the auditory domain. For example, it has been used to estimate the auditory filter shape (Shen and Richards, 2013, Shen et al., 2014, Schlittenlacher et al., 2020), the audiogram (Song et al., 2015, Cox and de Vries, 2021), the equal-loudness contour (Shen et al., 2018, Schlittenlacher and Moore, 2020, [Shen et al. 2024 JASA under-review]), and the band importance function for speech intelligibility (Shen and Kern, 2018 Jan-Dec, Shen et al., 2020, Shen and Langley, 2023).

Many of the previously mentioned studies use a parametric approach, which is suitable when there is sufficient *a priori* knowledge to formulate the model. In many cases, however, there is not sufficient information to formulate such a parametric model, and non-parametric modeling approaches may be preferred (Audiffren, 2022). A Gaussian Process (GP) model is a non-parametric, probabilistic alternative (Rasmussen and Nickisch, 2010). Gaussian process models assume that measurements follow a Gaussian distribution, described by a mean and variance. Rather than formulating how responses vary with test condition, GP models specify a set of kernel functions that govern how responses collected from a pair of test conditions may covary. This is considered non-parametric because how the measurement result varies with test condition is not explicitly formulated.

Several studies have paired a GP model with a Bayesian adaptive sampling strategy for rapid estimation of the audiogram (e.g., Song et al., 2015, 2018; Schlittenlacher et al., 2018; Cox and de Vries, 2021). In Song et al. (2015), the stimuli consisted of pure-tone frequencies presented at different intensities. The input to the GP model was frequency and intensity of the presented stimuli, and the listeners response of having heard or not heard the tone. The model is trained by the data to predict a listeners response across a wide range of frequencies and intensities. A linear kernel applied to the intensity dimension imposes monotonic growth in probability of tone detection with tone intensity, for a given frequency. A squared exponential kernel applied across the frequency dimension imposes that the threshold must be continuous. Model predictions are used to adaptively choose stimuli that are close to the threshold contour, thereby maximizing information gain. Besides efficiency, one advantage of this approach afforded by the GP model is a threshold contour that is continuous in frequency, rather than at discrete frequencies typical of traditional audiograms.

Schlittenlacher et al. (2018) also paired a GP model with a Bayesian adaptive sampling strategy for rapid estimation of the audiogram. The first approach in this study presented six tones at a given frequency with decreasing intensities, and the task of the listener is to count the number of tones heard. The adaptive sampling strategy placed the intensity range of these tones close to the estimated threshold at each iteration, thereby maximizing information gain. The second approach in this study is similar to Song et al. (2015), but with an extension of the GP model to account for lapses in attention. Both methods provided threshold estimates in approximately 4 minutes that match audiograms obtained using clinical best practice.

A Gaussian process model may also be well-suited for modeling ABR waveforms and the estimation of tone-burst ABR thresholds. Because it is a non-parametric model, it is able to characterize a large range of waveform morphologies, for which an explicit formulation may not be possible. This also makes a GP model well-suited for use across species and different hearing pathologies. Additionally, a GP model is able to leverage the covariance between waveforms in the time, frequency, and level dimensions simultaneously. The current study pairs a GP model with a Bayesian adaptive stimulus selection scheme for rapid estimation of tone-burst ABR thresholds across a wide frequency range.

The algorithm is implemented in a *post-hoc* numerical experiment using previously collected ABR datasets from four species of animals across five different laboratories. The datasets comprise averaged tone-burst ABR waveforms at systematically varying frequency and level conditions. From the full datasets, the algorithm utilizes a subset of stimuli conditions that are selected by the adaptive algorithm. The estimates using the adaptive algorithm were compared against threshold estimates by human raters who visually inspected the full data set. If close agreement between the adaptive algorithm and the human rater estimated threshold is observed, then it would indicate that the adaptive algorithm may be useful as a significant time-saving tool in a laboratory setting.

In what follows, Sect. 2 describes the implementation of the algorithm including the Gaussian process model and adaptive stimulus selection. The simulated experiment, including a description of the datasets is provided in Section 3. The results and discussion are in Sections 4 and 5, respectively.

## 2. Description of the algorithm

The adaptive algorithm leverages a GP model to predict the tone-burst ABR waveforms over a wide range of frequency and level combinations. During the procedure, the GP model is fitted to the tone-burst ABR waveforms collected for all previously tested stimulus conditions. The fitted model is then used to optimize the choice of the subsequent stimulus to minimize the expected uncertainty of the model predicted ABR threshold across frequencies. See Fig. 1 for a schematic of the algorithm.

**Figure 1:**
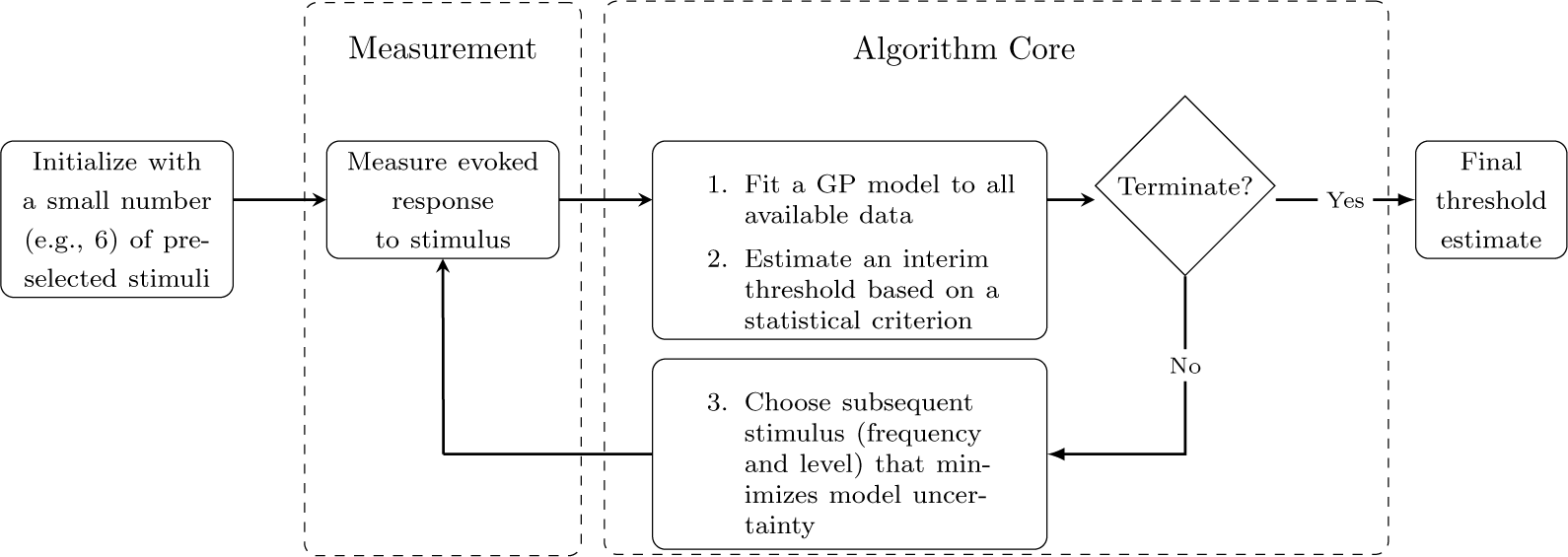
Schematic of the Bayesian adaptive algorithm for estimating hearing threshold. The adaptive algorithm intelligently samples the stimulus space by prioritizing the most informative stimulus conditions.

### 2.1. Gaussian process model

A Gaussian process model is a non-parametric, probabilistic model that relies on kernels that encode information about the covariance between data points across the stimulus space. A GP model is typically trained on data from a subset of the parameter space, and is used to make predictions across the entirety of the parameter space. Kernels may be chosen based on expected characteristics of the data.

The current algorithm fits a GP model (see Rasmussen and Nickisch, 2010) to the time-domain tone-burst ABR waveforms measured at certain frequencies and levels. The GP model imposes covariance kernels across three dimensions: time, frequency, and stimulus level, which are fitted simultaneously by the GP model. The fitted GP model is then used to predict the expected waveforms at other frequency and level combinations.

The GP model describes the covariance between random pairs of data points. For a given stimulus dimension (e.g., time), the ABR amplitude measured at two separate points (e.g., two points in time) along the dimension are *x_p_* and *x_q_*. The kernel *k*(*x_p_, x_q_*) describes the covariance between the ABR amplitudes measured at these two points. Kernels can be combined to form more complex composite kernels based on the needs of the model. Two types of kernels are used in the current GP model implementation (Rasmussen and Nickisch, 2010): the linear kernel with isotropic distance measure (covLINiso)

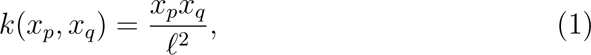

and the squared exponential with isotropic distance measure (covSEiso)

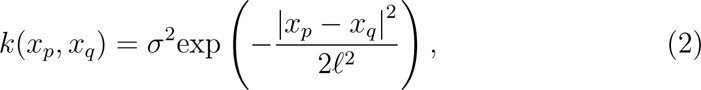

where *ℓ* is a characteristic length scale and *σ* is the signal variance. The linear kernel has one hyperparameter: hyp = [log (*ℓ*)]; the squared exponential has two: hyp = [log (*ℓ*); log (*σ*)]. Note that the characteristic length scale *ℓ* may have a different value in each kernel.

The kernels are chosen based on *a priori* expectations of the data covariance across the dimensions of time, frequency, and level. The squared exponential kernel is used for the time dimension because it is assumed data that are closer together in time should covary more than data that are far apart. Similarly, the squared exponential is chosen for the base-two logarithm of the frequency dimension, under the assumption that nearby frequency bands should covary more than ones that are far apart. This has the added consequence of assuring a threshold contour that is continuous across frequency and that exhibits a certain level of smoothness.

The covariance kernel characterizing the stimulus level is the product of the squared exponential and linear kernels. Pilot studies used the linear kernel alone, under the assumption that the waveform response amplitude should be proportional to the stimulus level. However, while linear amplitude growth with stimulus level is a reasonable characterization of ABR waveforms for some levels above threshold, it does not hold true below threshold, where the waveform is dominated by the measurement noise and becomes independent of stimulus level. For this reason, imposing the linear kernel alone results in predicted waveforms at levels well below threshold. The composite kernel combining the squared exponential and linear kernels allows the predicted waveforms to drop to zero amplitude for stimuli below threshold.

A composite kernel with seven hyperparameters is defined by multiplying the kernels defined for the dimensions: time (kernel: squared exponential), frequency (kernel: squared exponential), and level (kernel: squared exponential and linear).

Fitting a GP model to a set of ABR data involves optimizing the hyperparameters of the kernels. The optimization process for each hyperparameter begins by randomly drawing an initial parameter value from a preset search range. The search ranges for the seven hyperparameters used in the current study are listed in Table 3. The search ranges from which initial hyperparameter values are drawn can be selected generally to work for a large number of animal species, but may also be specifically tailored for a given specie based on previous research. The effect of hyperparameter initialization is discussed in Sect. 5.3.2.

Once the hyperparameters are fitted to the data, the GP model can be used to predict the tone-burst ABR waveforms across a desired range of frequencies and levels. The model predicted waveforms consist of both the mean and variance of the amplitude. The value of the variance can be interpreted as the uncertainty in the model with regard to the ABR amplitude for a given time, frequency, and level.

### 2.2. Adaptive algorithm

The adaptive algorithm uses an initial pre-selected set of test conditions (Sec. 2.2.1). Once the ABR waveforms for these initial test conditions are obtained, the iterative process involves three steps. First (#1 in Fig. 1), the GP model is fitted to all ABR waveforms collected so far. Second, (#2 in Fig. 1), an interim threshold is estimated (Sec. 2.2.2). Third (#3 in Fig. 1), a new test stimulus is determined (Sec. 2.2.3).

#### 2.2.1. Initial stimuli

A small set of initial stimuli (e.g., 6) are used to obtain the first set of tone-burst ABR waveforms used to fit the GP model. These stimuli are chosen at predominantly high levels in order to ensure that the GP model can correctly characterize the waveform morphology and dependence of the peak-to-peak amplitude on stimulus level. Provided these two characteristics are captured, the specific choice of initial stimuli does not appear to have a significant impact on the final threshold estimate.

#### 2.2.2. Threshold estimate

The threshold is estimated from the predicted tone-burst ABR waveforms returned by the GP model. The threshold at each frequency is defined as the lowest stimulus level for which the model predicted peak-to-peak amplitude of the tone-burst ABR waveform exceeds a given tolerance. This tolerance may be set as a constant or set based on the relative strength of the brainstem response to the measurement noise. A higher tolerance leads to higher predicted thresholds, while a lower tolerance leads to lower threshold estimates, see Sect. 5.3.1.

#### 2.2.3. Choice of subsequent stimulus

The subsequent stimulus is chosen at the level and frequency along the estimated threshold contour that has the largest variance value. This may be interpreted as sampling at the point along the estimated threshold contour for which the GP model is the least confident.

## 3. Experimental procedures

The current study aimed to provide initial validations of the adaptive algorithm for ABR threshold estimation through numerical simulations. The simulation experiment was based on five previously collected datasets from various animal species, collected by different laboratories. Each dataset consisted of tone-burst ABR waveforms for a pre-defined list of stimuli determined by the original research goals during data collection. See Sect. 3.2 for dataset specific information.

The current approach was to simulate novel runs of the adaptive algorithm by subsampling these existing datasets. It was expected that the adaptive algorithm could achieve comparable threshold estimates with respect to the human rater estimated threshold with much reduced testing time.

### 3.1. Simulated ABR

During each iteration of a simulated run of the adaptive procedure, a tone-burst ABR waveform for each level and frequency was simulated as the sum of the measured averaged tone-burst ABR waveform drawn from the dataset for the same level and frequency and a simulated noise. The noise was assumed to follow a Gaussian probability distribution that has been filtered and normalized by values obtained from measurements. It was simulated by: (1) drawing a vector of normally distributed random numbers; (2) lowpass filtering (6th-order Butterworth) this vector at the cutoff frequency used during the original data collection; (3) normalizing the amplitude of the filtered vector to obtain a root-mean-square (RMS) value comparable to a 6 ms section of the original data where no neural response is present. The RMS value of the waveform was calculated in the time-domain and the cutoff frequency of the low-pass filter applied during data collection was determined by visual inspection of the spectrum, if it was not available in the dataset metadata (see Table 1).

**Table 1:** An overview of the datasets includes the per-ear number of stimuli N_stim_, frequencies N_freqs_, averages N_avg_, and the root-mean-square estimate of measurement noise *x*_rms_ (*µ*V), averaged across all frequencies and levels for a given dataset. M1, M2, B1, G1, and GP1 refer to the first Mouse, second Mouse, Budgerigar, Gerbil, and Guinea Pig datasets, respectively.

| Dataset | $N_{\text{stim}}$ | $N_{\text{freqs}}$ | $N_{\text{avg}}$ | $x_{\text{rms}} (\mu\text{V})$ |
| --- | --- | --- | --- | --- |
| M1 | 102-106 | 7 | <256 | 0.12 |
| M2 | 68-85 | 4-5 | 300 | 0.10 |
| B1 | 72 | 6 | 300 | 0.42 |
| G1 | 66-114 | 6 | 512 | 0.14 |
| GP1 | 69-70 | 6 | 512 | 0.08 |

The noise was meant to replicate realistic conditions in which, were a measurement repeated at the same frequency and level, the simulated toneburst ABR waveform would include statistical variation matching the noise in the original measurement conditions. Therefore, the RMS scaling factor was different for each stimulus condition, based on the RMS noise present in the original data. Additionally, an independent sample of noise was drawn from the distribution for each stimulus condition, including the case in which the same stimulus was chosen more than once. See Fig. 2 for an example of a measured and simulated waveform.

**Figure 2:**
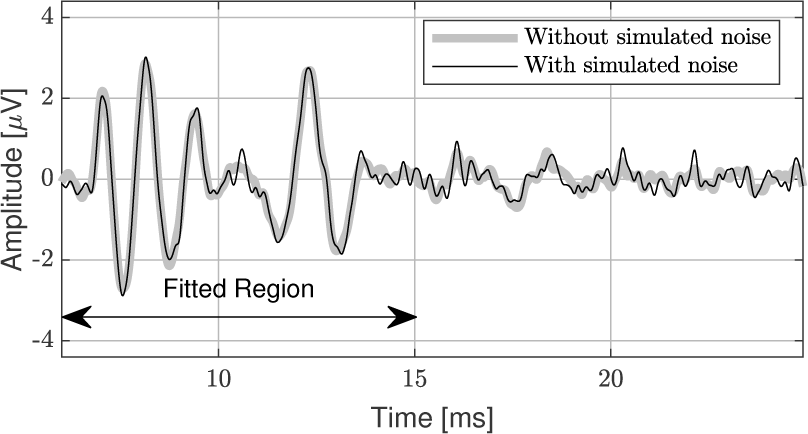
Measured waveform (M1E1, 4 kHz at 80 dB SPL) with and without additional simulated noise.

### 3.2. Datasets

The algorithm was tested in a simulated experiment using five existing ABR datasets provided by different laboratories. The tone-burst ABR waveforms in each dataset consist of time-synchronously averaged measurements. The specific parameters of the datasets, such as the number of averages, sampling rate, and stimuli conditions, differ due to the varying requirements of the research for which they were collected. A summary for each is provided below as well as in Table 1.

#### 3.2.1. Dataset 1: Mouse (M1)

The first dataset consists of tone-burst ABR waveforms from two Normal-Hearing (NH) mice ears (unpublished data, collected following a similar ABR protocol as specified in Tan et al., 2020). The data is collected for tone-burst stimuli at 4000, 5656, 8000, 11313, 16000, 22627, 32000 Hz, at stimulus presentation levels ranging from 80 dB SPL in 5 dB decrements, down to 55 dB SPL at 4000 Hz, 40 dB SPL at 32000 Hz, and 5 dB SPL for all other frequencies. The time-synchronously averaged waveforms for each stimulus condition consists of up to a maximum of 264 averages, but often has fewer than 100 averages for the highest stimuli levels. The dataset provided approximately 100 total stimuli conditions for each ear.

#### 3.2.2. Dataset 2: Mouse (M2)

The second dataset consists of ABR recordings from four NH and five Hearing-impaired (HI) mice, for a total of 9 unique ears, collected in a protocol similar to Zhang et al., 2022. The HI ears were exposed to Kainic-acid. The ABR waveforms were collected for tone-burst stimuli at 4000, 8000, 16000, 24000, and 32000 Hz for the NH ears, and for 8000, 16000, 24000, and 32000 Hz for the HI ears. Stimuli presentation levels range from 10-90 dB SPL in 5 dB increments. The time-synchronously averaged waveforms for each stimulus condition consists of 300 averages. The dataset provided 85 total stimuli conditions for each pre-exposure ear and 68 for each post-exposure ear.

#### 3.2.3. Dataset 3: Budgerigar (B1)

The third dataset consists of ABR recordings from four Budgerigar ears pre- and post-Kainic-acid exposure (Henry and Abrams, 2018). The ABR waveforms were collected using tone-burst stimuli at 500, 1000, 2000, 3000, and 4000 Hz with levels ranging from 15 to 80 dB SPL in 5 dB increments, skipping the 65 and 75 dB SPL levels. The time-synchronously averaged waveforms for each stimulus condition consists of 300 averages. The dataset provided 72 total stimuli conditions for each ear.

#### 3.2.4. Dataset 4: Gerbil (G1)

The fourth dataset consists of baseline ABR recordings from three Gerbil ears and recordings for two Gerbils 42 days post cochlear implantation (Private correspondence, from the lab of Christoph Arnoldner, Department of Otorhinolaryngolog, Medical University of Vienna). The data is collected at 1, 2, 4, 8, 16, and 32 kHz, with levels typically ranging from 15 to 65 dB SPL in 5 dB steps for the baseline data, and 50-110 dB SPL for post cochlear implantation. The time-synchronously averaged waveforms for each stimulus condition consists of 512 averages. The dataset provided 66 stimuli conditions for the baseline data, with 78 and 114 stimuli conditions for the two post-cochlear implantation data.

#### 3.2.5. Dataset 5: Guinea Pig (GP1)

The fifth dataset consists of baseline ABR recordings for three Guinea Pig ears (Private correspondence, from the lab of Christoph Arnoldner, Department of Otorhinolaryngolog, Medical University of Vienna). The data is collected at 1, 2, 4, 8, 16, and 32 kHz, ranging from 15 to 75 dB SPL for ear GP1E1, 25 to 75 dB SPL for ear GP1E2 (missing 65 dB SPL tone at 4 kHz), and 15 to 65 dB SPL for ear GP1E3, in 5 dB increments. The time-synchronously averaged waveforms for each stimulus condition consists of 512 averages. The dataset provided approximately 70 stimuli per ear.

### 3.3. Algorithm configurations

#### 3.3.1. Data preprocessing

For each dataset, the magnitude of the waveforms was normalized so that each dataset was expressed in microvolts. The tone-burst ABR waveforms were windowed to a 10 ms (see Fig. 2) frame beginning at 5 ms after stimulus onset. This window encompassed the primary evoked potential waveform features, e.g., ABR wave III and V. The data were all re-sampled to a common sampling rate of 6 kHz, which is sufficiently high to capture the main waveform characteristics. This sampling rate ensures that the computation time of fitting the GP model remains between 15-80 s, depending on the number of stimuli.

#### 3.3.2. Tolerance for threshold

The minimum peak-to-peak amplitude, above which a waveform is deemed present, was determined during pilot simulations. Each dataset requires a unique tolerance, reflecting the differences data collection methods (e.g., number of averages, background noise, etc.) as well as variation in waveform morphology. The same tolerance is used for each ear in a given dataset. Tolerances were set to 0.68, 0.44, 2.00, 0.66, and 0.70 *µV*, for the M1, M2, B1, G1, and GP1 datasets, respectively. In general, a smaller tolerance will result in a lower threshold prediction for a given dataset, while a larger tolerance will result in a higher threshold prediction, see Sect. 5.3.1.

#### 3.3.3. Initialization stimuli

The frequencies and levels of the initialization stimuli for each dataset are summarized in Table 2. The initialization stimuli are different for each dataset, and occasionally within datasets, reflecting the inhomogeneity of the stimulus space chosen by the laboratories that provided the data for this algorithm development. Initialization stimuli are chosen to include the highest level for 4 frequencies spanning the available frequency range. Additionally, levels 20 and 30 dB below the highest level are chosen for two frequencies near the middle of the available frequency range.

**Table 2:** Initialization stimuli for each dataset: Mouse (M1 and M2), Budgerigar (B1), Gerbil (G1), and Guinea Pig (GP1). For each dataset, initialization stimuli are chosen to include the highest level for a wide range of frequencies, and levels 20-30 dB below the highest level for some middle frequencies. As long as this general scheme is used, the performance of the algorithm is not very sensitive to the specific combination of initialization stimuli. (*) Ears M2E5-M2E9 do not have data for 4.0 kHz; for these ears the initial stimulus of 90 dB SPL is included at for 8.0 kHz, rather than 4.0 kHz. (**) The post cochlear implanted ears G1E1: post-exp. and G1E3: post-exp. were tested at higher levels, up to a maximum of 110 dB SPL. For these ears, the same scheme is used, but shifted up from a maximum level of 65 dB SPL to a maximum level of 110 dB SPL. (* * *) Ear GP1E3 was tested at lower levels, up to a maximum of 65 dB SPL. For this ear, the same scheme is used, but shifted down from a maximum level of 75 dB SPL to a maximum level of 65 dB SPL.

| Dataset | Frequency<br>kHz | Initialization<br>Level dB SPL |
| --- | --- | --- |
| M1 | 4.0 | 80 |
|  | 5.7 | - |
|  | 8.0 | 80, 60 |
|  | 11.3 | - |
|  | 16.0 | 80, 50 |
|  | 22.6 | - |
|  | 32 | 80 |
| M2 * | 4.0 | 90 |
|  | 8.0 | - |
|  | 16.0 | 90, 70 |
|  | 24.0 | 90, 60 |
|  | 32.0 | 90 |
| B1 | 0.5 | 80 |
|  | 1.0 | - |
|  | 2.0 | 80, 60 |
|  | 3.0 | 80, 50 |
|  | 4.0 | - |
|  | 6.0 | 80 |
| G1 ** | 1.0 | 65 |
|  | 2.0 | - |
|  | 4.0 | 65, 45 |
|  | 8.0 | 65, 35 |
|  | 16.0 | - |
|  | 32.0 | 65 |
| GP1 * * * | 1.0 | 75 |
|  | 2.0 | - |
|  | 4.0 | 75, 55 |
|  | 8.0 | 75, 45 |
|  | 16.0 | - |
|  | 32.0 | 75 |

The rational for using the highest level for many of the initialization stimuli is to ensue that the GP model is provided with a clear example of the tone-burst ABR waveform. The two stimuli at 20 to 30 dB below the highest level are chosen because they typically still contain a discernible ABR, but at a lower amplitude than the highest levels: providing the GP model with data that reinforces the predicted behavior that ABR amplitude should increase with the level. In practice, the specific collection of initialization stimuli does not have a large impact on the performance of the algorithm, as long as some data with strong ABRs are included.

#### 3.3.4. Hyperparameter initialization

The initialization range of the hyperparameters are summarized in Table 3. The ranges work well for the adaptive procedure for all datasets. Identifying appropriate values may depend on the dataset parameters, especially the sampling frequency, which can vary by orders of magnitude between experimental set-ups. See Sect. 5.3.2 for a discussion about hyperparameter initialization schemes.

**Table 3:** Range from which each hyperparameter is randomly drawn for M1, M2, G1, and GP1 datasets. Dataset B1 is the same, except the time-domain hyperparameter log (*ℓ*) is drawn from [*−*12*, −*10].

| time |  | level |  |  | frequency |  |
| --- | --- | --- | --- | --- | --- | --- |
| sq. exp. |  | lin. | sq. exp. |  | sq. exp. |  |
| $\log(\ell)$ | $\sigma$ | $\log(\ell)$ | $\log(\ell)$ | $\sigma$ | $\log(\ell)$ | $\sigma$ |
| -10, -8 | -5, -1 | 2, 4 | 2, 4 | 1, 2 | -2, 0 | 1, 3 |

#### 3.3.5. Ensuring an acceptable GP model fit

An acceptable GP model fit is required before a subsequent stimulus is selected. Occasionally, a failed GP model fit will return predicted tone-burst ABR waveforms with nearly zero amplitude (less than e-10 V) for all stimuli conditions, indicating a failed model fit. Therefore, after each fit, it is verified that there is at least one predicted waveform with a non-zero amplitude. In this experiment, a peak-to-peak amplitude of less than 9 *nV* is considered to have zero amplitude. This minimum amplitude requirement is considerably smaller than the ABR response (approximately 5 *µV*) and background noise (1 *µV*) in the measurements.

When the GP model fails to achieve a satisfactory fit, another attempted fit is made using a new set of hyperparameter initialization values drawn from the ranges described in Sect. 3.3.4. If the GP model fails to fit 10 consecutive times, the current waveform is discarded and a new stimulus is drawn from a random frequency at the highest level available in the dataset. The rational for choosing a high level is to provide the GP model with a waveform that has the highest likelihood of containing a large ABR peak, increasing the likelihood of a successful model fit.

Note that, when a waveform is discarded due to an unsatisfactory GP model fit, the stimulus condition (frequency and level) is retained in available dataset. That is, the same stimulus condition could be chosen by the algorithm in a later iteration. However, a subsequently drawn waveform at the same stimulus condition will have a new sample of noise added to the measurement as described in Sect. 3.1.

#### 3.3.6. Termination criteria

The algorithm was run until a fixed number of stimuli were included in the subset of the stimulus space. In the current study, all results are shown for a fixed number of 20 stimuli, see Sect. 5.3.3.

### 3.4. Evaluation of algorithm

The algorithm is validated with respect to human rater estimates of threshold. For each ear, four consecutive runs of the algorithm provides four threshold estimates. Independently, four human raters estimate threshold (see Section 5.2.2). The mean of the threshold estimates from the four consecutive runs of the algorithm is compared against the mean of the threshold estimates four human raters to evaluate general agreement between the algorithm and human rater estimates. Estimated thresholds within 10 dB are considered satisfactory.

Additionally, the standard deviation of the adaptive algorithm threshold estimates are compared with the standard deviation of the human rater estimates. A smaller standard deviation of the algorithm estimates indicates that the test/retest reliability of the algorithm threshold estimate is better than the the inter-rater consistency for the human raters, and vice versa.

#### 3.4.1. Human raters

The four human raters estimated threshold by visual inspection of the tone-burst ABR waveforms for all levels available in the dataset one frequency at a time, with no indication of dataset specifics such as species, expected sensitivity, or expected waveform morphology. Instructions were given to identify the ABR response at the highest stimulus level and visually track the response to lower levels. The lowest level at which a waveform can still be identified is the human estimated threshold. The raters all have some practice inspecting ABR waveforms, and one has clinical training as an audiologist.

## 4. Results

The mean thresholds are plotted in Figs. 3-7, where the four adaptive algorithm estimates are denoted by the left pointing blue triangles (color online). The error bars depict the standard deviations. Similarly, the mean and standard deviation of the four human rater estimates are shown by the right pointing red triangles. The annotation of form m/n shows m, the number of waveforms used in the algorithm, and n, the total number of waveforms available in the dataset viewed by the human raters.

**Figure 3:**
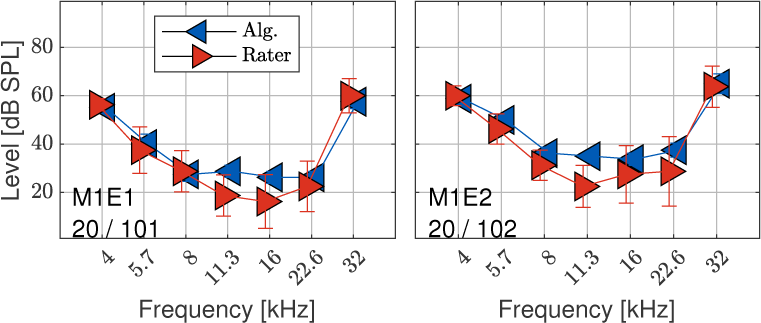
Mouse set 1: the mean threshold estimate from 4 human raters and 4 successive runs of the algorithm are shown by the right and left pointing triangle data makers, respectively (color online). Error bars denote the standard deviation. Data markers and error bars offset horizontally for legibility.

Two types of single-value metrics are used to compare the mean algorithm estimated threshold with the mean human rater estimated threshold. The first is the *absolute difference*, *ɛ*_abs_, which is the magnitude of the difference in mean threshold estimates averaged over frequency. The second is the *bias difference*, *ɛ*_bias_, which is the (signed) difference in mean threshold estimates averaged over frequency.

Additionally, a metric relating to the variability of estimates for a given ear is calculated as the standard deviation of threshold estimates averaged over frequency, 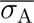 and 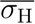, where subscripts A and H refer to the algorithm and human rater estimates, respectively.

The outcomes of these four metrics (*ɛ*_abs_*, ɛ*_bias_, 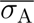 and 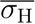) are summarized in Table 4 for each ear.

**Table 4:** Summary of the absolute *ɛ*_abs_ and bias *ɛ*_bias_ of the adaptive algorithm estimated threshold relative to human estimated threshold, averaged across frequencies; standard deviation of the four consecutive adaptive algorithm runs 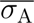 and the standard deviation of the four human raters 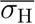, averaged across frequencies. All values in dB.

| ear | $\epsilon_{\text{abs}}$ | $\epsilon_{\text{bias}}$ | $\overline{\sigma}_{\text{H}}$ | $\overline{\sigma}_{\text{A}}$ |
| --- | --- | --- | --- | --- |
| M1E1 | 4 | -3 | 8 | 2 |
| M1E2 | 5 | -5 | 8 | 2 |
| M2E1 (pre) | 3 | 3 | 11 | 6 |
| M2E2 (pre) | 4 | 4 | 13 | 2 |
| M2E3 (pre) | 22 | -22 | 11 | 1 |
| M2E4 (pre) | 8 | 5 | 14 | 3 |
| M2E5 (post) | 7 | -6 | 13 | 2 |
| M2E6 (post) | 8 | -8 | 9 | 3 |
| M2E7 (post) | 14 | -1 | 9 | 4 |
| M2E8 (post) | 6 | 3 | 18 | 4 |
| M2E9 (post) | 0 | 0 | 0 | 0 |
| B1E1 (pre) | 9 | 3 | 9 | 4 |
| B1E1 (post) | 5 | -5 | 7 | 1 |
| B1E2 (pre) | 2 | -0 | 15 | 3 |
| B1E2 (post) | 11 | -11 | 13 | 2 |
| B1E3 (pre) | 6 | 2 | 14 | 4 |
| B1E3 (post) | 9 | 1 | 17 | 10 |
| B1E4 (pre) | 10 | -10 | 13 | 2 |
| B1E4 (post) | 14 | -14 | 11 | 3 |
| G1E1 (pre) | 12 | -9 | 11 | 5 |
| G1E1 (post) | 9 | 9 | 19 | 5 |
| G1E2 (pre) | 6 | 5 | 7 | 6 |
| G1E3 (pre) | 8 | 8 | 17 | 2 |
| G1E3 (post) | 14 | 14 | 18 | 8 |
| GP1E1 | 5 | 1 | 13 | 5 |
| GP1E2 | 13 | -11 | 13 | 6 |
| GP1E3 | 14 | -11 | 14 | 4 |

### 4.1. Mouse 1

The threshold estimates for the Mouse 1 dataset are plotted in Fig. 3, where the left and right panels correspond to two different ears. The absolute and bias difference metrics, *ɛ*_abs_ and *ɛ*_bias_, are less than 10 dB for two out of two ears (see Table 4), indicating satisfactory performance per ear, and on the dataset as a whole. The bias difference is negative for both ears, with a mean value of -4.1 dB, indicating that the tolerance defined for threshold (peak-to-peak amplitude) used by the algorithm may be somewhat more conservative than human raters.

The standard deviation metric for algorithm performance 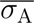 is less than 10 dB for nine out of nine ears (dataset mean: 2.3 dB), while that of the human raters is less than 10 dB for three out of nine ears (dataset mean: 8.4 dB). This indicates that the variability of threshold estimates from one run of the adaptive procedure to another is smaller than the variability across individual, non-expert human raters.

### 4.2. Mouse 2

The threshold estimates for the Mouse 2 dataset are plotted in Fig. 3, where the four ears without Kainic-acid exposure are shown in the left panels, and the five ears with Kainic-acid exposure in the right panels. The absolute difference is less than 10 dB for seven out of nine ears (exceptions: M2E3, M2E7), and the bias difference is less than 10 dB for eight out of nine ears (exception: M2E3), indicating satisfactory performance for the majority of the dataset.

The standard deviation metric for algorithm performance 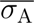 is less than 10 dB for nine out of nine ears (dataset mean: 3.1), while that of the human raters is less than 10 dB for three out of nine ears (dataset mean: 11.2). This indicates that the variability of threshold estimates from one run of the adaptive procedure to another is smaller than the variability across individual, non-expert human raters.

### 4.3. Budgerigar

The threshold estimates for the Budgerigar dataset are plotted in Fig. 5, where the left panels show the estimates pre Kainic-acid exposure and the right panels show the same ears post Kainic-acid exposure. The absolute and bias difference is less than 10 dB for five out of eight ears (exceptions: B1E2 post, B1E4 pre and post), indicating satisfactory performance for the majority of the dataset.

**Figure 4:**
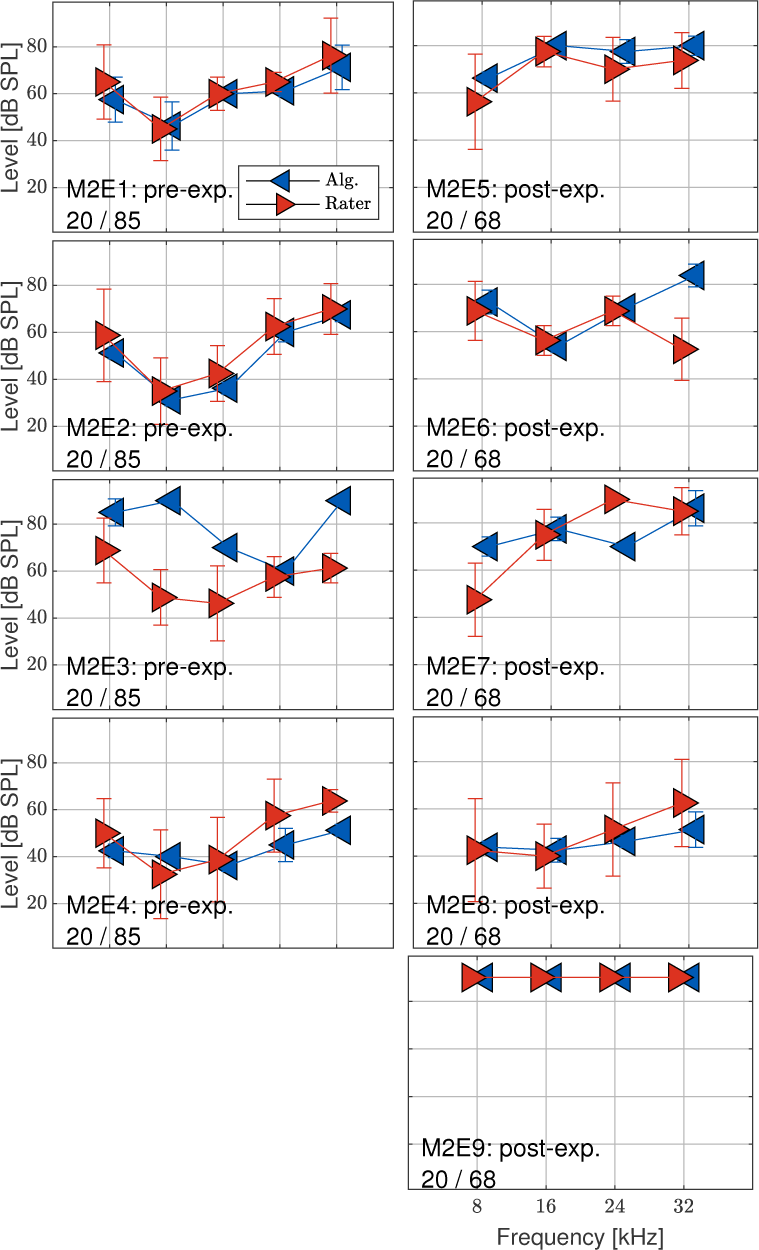
Mouse dataset 2: data markers are the same as in Fig. 3. Ears with and without Kainic-acid exposure are shown in the right and left panels, respectively.

**Figure 5:**
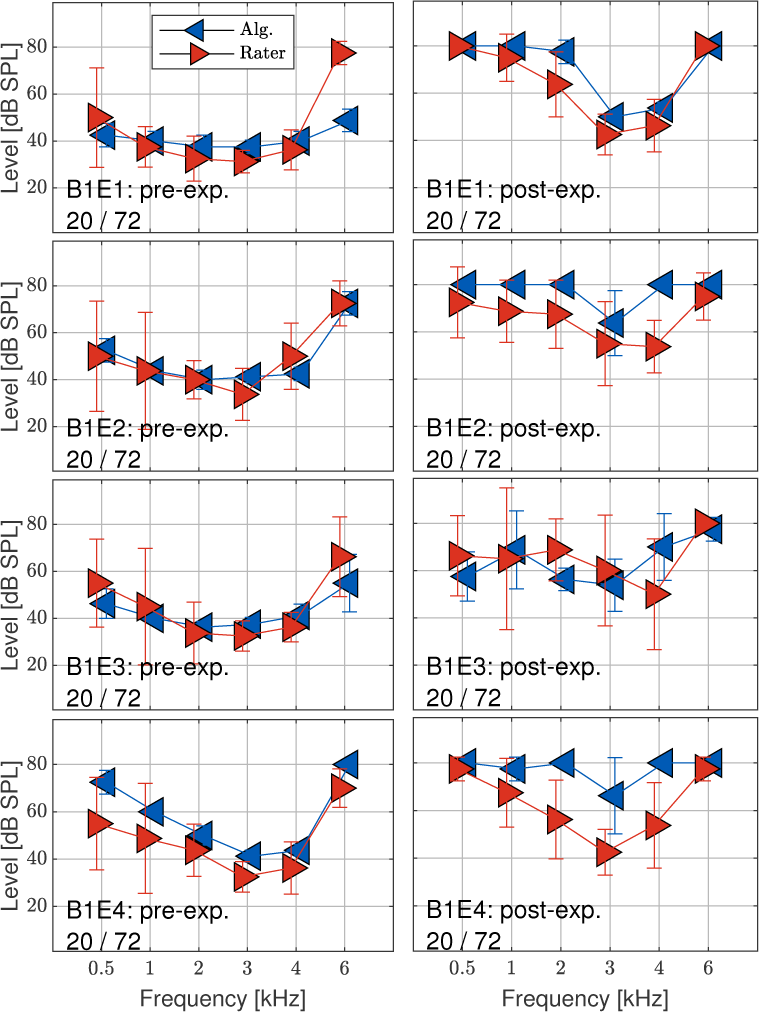
Budgerigar dataset: data markers are the same as in Fig. 3. Ears pre- and post-Kainic-acid exposure are shown in the left and right panels, respectively.

The standard deviation metric for algorithm performance 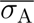 is less than or equal to 10 dB for eight out of eight ears (dataset mean: 4.1), while that of the human raters is less than 10 dB for two out of eight ears (dataset mean: 12.8). This indicates that the variability of threshold estimates from one run of the adaptive procedure to another is less than the variability across individual, non-expert human raters.

There are differences between how the algorithm performed on the pre- and post-exposed ears. The average absolute and bias difference metrics of the pre-exposed ears are *ɛ*_abs_ = 7.1 and *ɛ*_bias_ = *−*1.4, while the averages for the post-exposed ears are *ɛ*_abs_ = 10.3 and 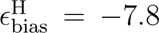. This shows that the algorithm estimated threshold matches the human rater estimates more closely and with less bias for the pre-exposed ears than the post-exposed ears.

The standard deviation metrics are the same (when rounded to the nearest integer) for the pre- and post-exposed ears: 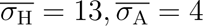.

### 4.4. Gerbil

The threshold estimates for the Gerbil dataset are plotted in Fig. 6, where the left panels show thresholds for ears pre-cochlear implantation and the right panels show thresholds post-cochlear implantation. The absolute difference is less than 10 dB for three out of five ears (exceptions: G1E1 pre, G1E3 post), while the bias difference is less than 10 dB for four out of five ears (exception: G1E3 post), indicating satisfactory performance for the majority of the ears in the dataset. However, the absolute and bias difference metrics averaged over the dataset are *ɛ*_abs_ = 10.2 and 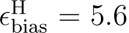, indicating that the algorithm did not have satisfactory performance with respect to the absolute difference on the dataset as a whole. Further, the bias difference metric indicate that tolerance defined for threshold (peak-to-peak amplitude) is generally less conservative compared to the human raters.

**Figure 6:**
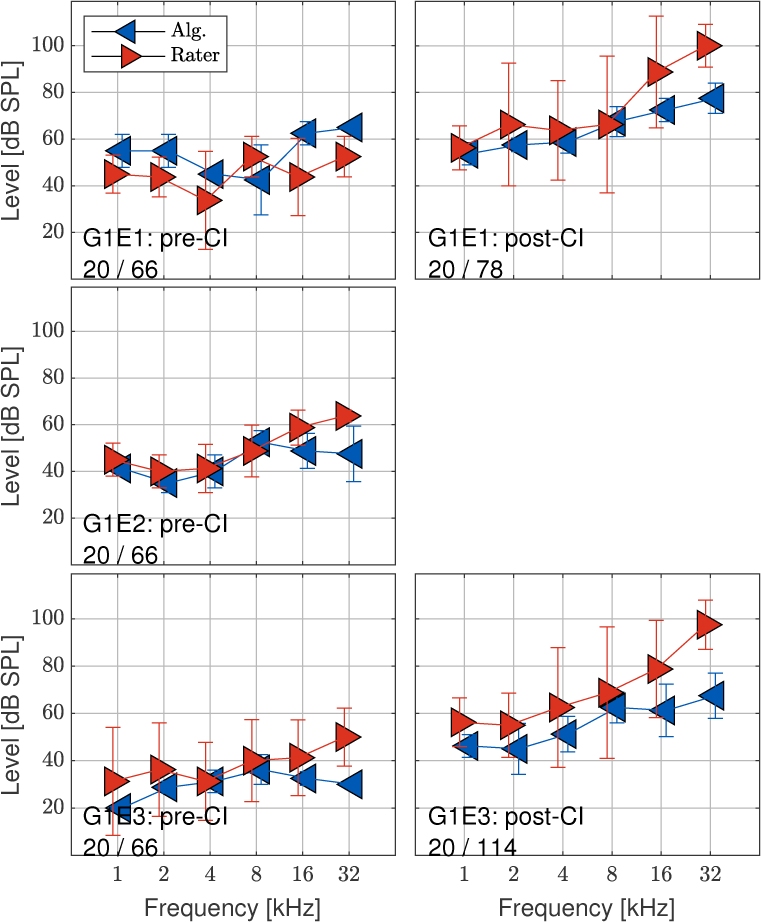
Gerbil dataset: data markers are the same as in Fig. 3. Left panels show baseline data and right panels show post cochlear implanted ears 1 and 3 (post-implantation data not available for ear 2).

The standard deviation metric for algorithm performance 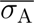 is less than 10 dB for five out of five ears (dataset mean: 5.6), while that of the human raters is less than 10 dB for one out of give ears (dataset mean: 15.0). This indicates that the variability of threshold estimates from one run of the adaptive procedure to another is less than the variability across individual, non-expert human raters.

The mean bias difference of the pre- and post-CI ears is 1.7 dB and 11.6 dB, respectively. The mean standard deviation for the pre-CI ears are 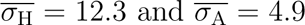, while the mean standard deviation for the post-CI ears are 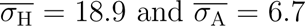. This reflects a greater ambiguity for both the human raters and adaptive algorithm while estimating the threshold contour of the post-implanted ears. Regardless, the algorithm has less variability compared to the human raters for both pre- and post-exposed ears.

### 4.5. Guinea Pig

The absolute and bias difference is less than 10 dB for one out of three ears (exceptions: GP1E2, GP1E3), indicating unsatisfactory performance for the majority of the ears in the dataset. However, there is a substantial mean bias difference is -7.2 dB, indicating that a less conservative threshold peak- to-peak tolerance value may result in smaller absolute and bias differences with respect to the human rates.

The standard deviation metric for algorithm performance 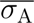 is less than 10 dB for three out of three ears (dataset mean: 5.6), while that of the human raters is less than 10 dB for zero out of three ears (dataset mean: 13.4). This indicates that the variability of threshold estimates from one run of the adaptive procedure to another is less than the variability across individual, non-expert human raters.

## 5. Discussion

The adaptive algorithm is able to estimate an ABR threshold with minimal customization across many different species and hearing pathologies. The adaptively estimated ABR thresholds match human rater estimates reasonably well (within 10 dB averaged over frequency) for 19 out of 27 ears. Systematic bias of threshold estimates in some datasets indicates that improved absolute difference values could be attained with different threshold tolerance values.

### 5.1. Relation to prior works

#### 5.1.1. Studies to improve ABR measurement efficiency

As discussed in Sect. 1, a number of approaches have been developed and successfully validated that reduce ABR measurement time by seeking to optimize the stimulus configuration. One such approach is to interleave frequencies, which allows for a higher stimulus presentation rate without neural adaptation (Buran et al., 2020). Another example is to adaptively choose the number of stimulus presentations used to generate the synchronously averaged waveform based on a signal-to-noise type metric: when sufficient signal has been detected in the averaged waveform, the protocol may advance to the subsequent stimulus (Elberling and Don, 1984). In both cases, the stimuli conditions of the measurements are typically pre-defined, so the total time-savings is proportional to the time-savings achieved for each stimulus condition.

The adaptive algorithm presented in the current article takes a different approach: rather than minimizing the time required to measure the ABR waveform for a given stimulus condition, time-savings is achieved by minimizing the total number of stimuli conditions that are necessary to estimate ABR threshold. These two approaches may be seen as complementary methods for minimizing ABR threshold estimation time.

#### 5.1.2. Studies to improve the efficiency of estimating behavioral thresholds

The use of GP models in the current article bears some similarities with their use in behavioral audiogram measurement methods (Song et al., 2015, 2018; Schlittenlacher et al., 2018; Cox and de Vries, 2021). All of these methods, including the adaptive algorithm in the current article, use a GP model to predict a response across a wide range of frequencies and levels, from which a continuous threshold can be defined. Additionally, all these methods impose basic assumptions on the nature of the threshold, e.g., continuity and a degree of smoothness across frequency, monotonic growth with intensity, and so forth. Further, the GP model is used not only for threshold prediction, but also stimulus optimization.

The GP model used in the other approaches rely on predicting the likelihood of a behavioral response yes/no for a given stimulus along a dimensionless scale of 0 to 1. This likelihood may be used to define a threshold; one such scheme is to define threshold at a certain response probability, e.g., 50%. In contrast, the proposed algorithm uses a GP model that performs regression rather than classification. Here the model operates along three dimensions (frequency, level, time) rather than two dimensions (frequency and level), and a threshold is defined not according to probability of a behavioral response, but instead from a minimum peak-to-peak amplitude of the predicted waveform. Therefore, the current algorithm has extended these previous efforts to the estimation of physiological-based threshold estimation.

### 5.2. Limitations

#### 5.2.1. Algorithm and model configurations

The algorithm presented in the current article has certain limitations that may be specific to its current configuration. The most consequential is that it was validated through numerical simulations using previously collected ABR datasets. It is not clear how the algorithm will generalize to *in situ* measurement protocols, especially in cases where the obtained data may differ from the currect datasets, e.g., for different species, differing hearing pathologies, stimulus parameters, etc. However, that the algorithm was successfully applied to 5 distinct datasets including 4 species through simulation provides support that it may be successful in a live experiment.

Another limitation is imposed by the set parameters of the previously collected datasets. Specifically, in each dataset, the stimuli range along systematic grids of previously defined frequencies and levels. Therefore, the adaptive algorithm is limited to choosing stimuli conditions that already exist in the datasets. Ideally, the algorithm should be able to specify any frequency or level, albeit constrained within pre-defined ranges for practical and safety considerations. Greater flexibility of stimulus parameter choice may improve the algorithm performance, as the optimal stimulus parameters may not exist in the five ABR datasets used in the current study. However, greater stimulus parameter flexibility may impact the optimal hyperparameter initialization and optimized values, which will have to be examined during an *in situ* validation of the algorithm.

Additionally, it may be noted that the authors make no claim that the algorithm is configured for maximal performance. Configurations such as the threshold definition, e.g., the peak-to-peak tolerance (Sect. 3.3.2), stimulus selection (2.2.3), hyperparameter initialization (Sect.3.3.4), the termination criteria (Sect.3.3.6), etc., may be improved upon or tailored for the needs of a specific ABR measurement protocol.

#### 5.2.2. Human raters

Another limitation with the current study is the considerable variation between the inter-rater estimated thresholds used as a reference against which to validate the adaptive algorithm threshold estimate. This variation may reflect that some raters are more conservative/permissive than others in defining threshold. Further, although the raters all have some experience in estimating ABR thresholds, none are experts for the species used in this article. More reliable ABR threshold estimates may lesson the inter-rater differences. Regardless, the same four raters were used throughout the study, so there is never the less some consistency for human-rater thresholds across datasets.

### 5.3. Effects of algorithm configuration

The development of the adaptive algorithm involved configurating parameters such as the threshold tolerance, hyperparameter initialization values, and termination criteria, based on preliminary numerical experiments using the available data. This section highlights some practical considerations to prepare the algorithm for novel experimental conditions.

#### 5.3.1. Threshold tolerance value

The threshold is identified as the lowest level for which the peak-to-peak amplitude of the predicted ABR waveform exceeds a given tolerance that is defined for each dataset (see Sect. 2.2.2). In general, a tolerance with a larger value reflects a more conservative criteria for the presence of an ABR waveform, which results in a higher threshold estimate. Conversely, a tolerance with a smaller value reflects a less conservative criteria and will typically result in a lower threshold estimate.

To demonstrate the effect of the tolerance value, threshold estimates using three different tolerance values are simulated for the M1E1 and M1E2 ears. The peak-to-peak tolerance values are set to 0.34, 0.68, and 1.02 *µV*, where 0.68 *µV* is the value used in the results presented in Sect. 4.1; 0.34 and 1.02 are 0.5 and 1.5 times this value, respectively. Figure 8 shows that a higher tolerance value typically results in higher threshold estimates.

**Figure 7:**
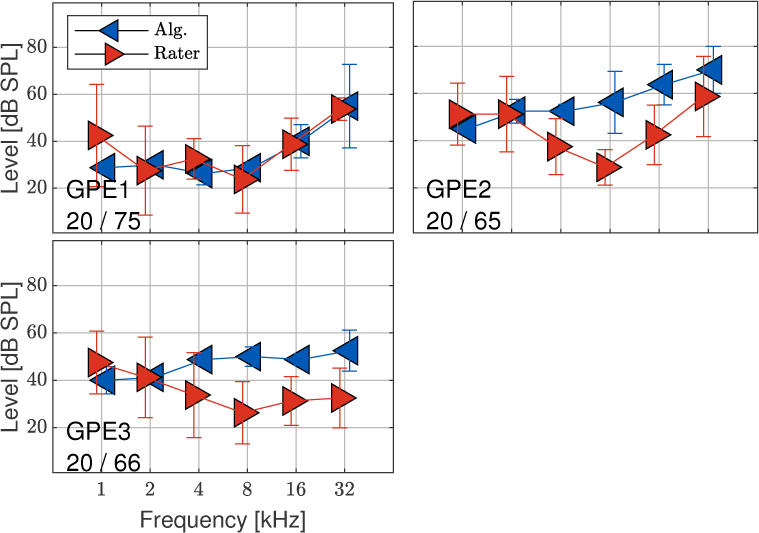
Guinea Pig dataset: data markers are the same as in Fig. 3.

**Figure 8:**
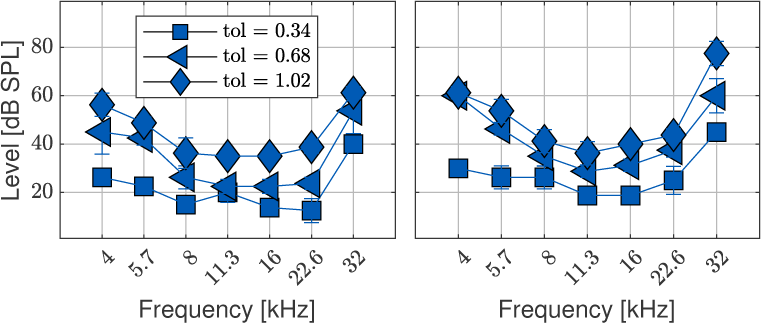
Threshold estimates for ears M1E1 and M1E2 using three tolerances values: 0.34, 0.68, and 1.02 are denoted by the square, triangle, and diamond markers, respectively. As in Figs. 3-7, the data markers and error bars show the mean and standard deviation of four independent simulated runs of the algorithm.

The choice of tolerance value must be tailored for a given experimental set-up (e.g., data acquisition hardware) and depends on overall peak-to-peak amplitude of the ABR waveforms, the amount of measurement noise, and how conservative an estimate is desired by the user. See Sect. 5.3.1 for a summary of the values used in the current article.

#### 5.3.2. Effect of hyperparameter initialization

The ranges of hyperparameter used to fit the GP model may impact the performance of the algorithm. These hyperparameters correspond to the covariance between data points along each of the three dimensions of the dataset: time, frequency of the stimulus, and intensity of the stimulus. Therefore suitable values depend on the discretization scheme along these dimensions, e.g., the sampling rate, base-two logarithm of the discrete stimulus frequencies, and inter-level gap between stimulus intensities.

The initialization ranges used for the adaptive algorithm in the current article (see Sect. 3.3.4 and Table 3) are the same across all five datasets (except the time-dimension characteristic length scale *ell* for dataset B1), indicating that they are likely a good starting point when applying this algorithm to novel datasets. However, these datasets have relatively similar discretization schemes: novel datasets that use very different discretization schemes, e.g., a much higher sampling rate, or denser set of stimulus frequencies or intensities, may need to modify the initialization ranges.

As an example, the length scale *ℓ* in the squared exponential kernel imposed on the frequency dimension determines the degree of covariance across frequencies and, in effect, governs how far away in the base-two logarithm of the frequency dimension the GP model is able to use information from one frequency to inform the predicted waveform at another frequency. A very small value of *ℓ* implies limited covariance across frequencies, while a large *ℓ* implies considerable covariance across frequencies.

To demonstrate the effect of the hyperparameter initialization ranges, a series of additional simulations are performed on the M1E1 ear. The initialization value of each parameter is fixed to the average value of the ranges provided for M1 in Table 3, except the length scale *ℓ* in the squared exponential kernel applied to the frequency domain, which is varied from log_2_(*ℓ*) = -4 to log_2_(*ℓ*) = 10. Simulations are repeated 10 times for each condition.

The top panel of Fig. 9 shows the optimized value of log_2_(*ℓ*) as a function of the initialization value. The line and error bars correspond to the mean and standard deviation of the 10 consecutive runs of the algorithm. For the lowest and highest initialization values, log_2_(*ℓ*) = -4 and log_2_(*ℓ*) = 10, the GP model fails to optimize this hyperparameter and the hyperparameter is left at its initialization value. For the lowest value, log_2_(*ℓ*) = -4, the GP model often fails to fit a stimulus condition, and in each run the simulations were terminated early after repeated failure to achieve a satisfactory model fit, see Sect. 3.3.5. At intermediate initialization values, the optimized values are all around log_2_(*ℓ*) = -1, irrespective of the initialization value.

**Figure 9:**
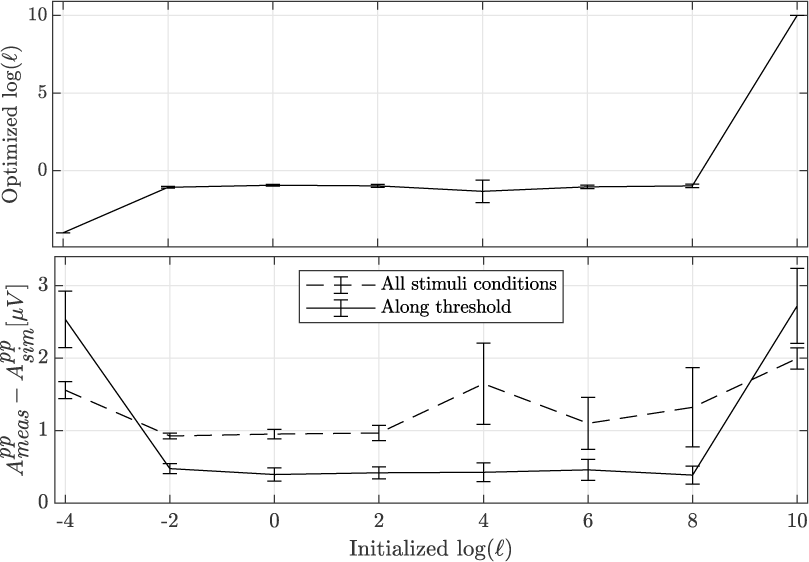
Both panels simulated for ear M1E1. Top panel: optimized log_2_(*ℓ*) as a function of initialization value of log_2_(*ℓ*) for the squared exponential kernel imposed on the frequency dimension. The line and error bars correspond to the mean and standard deviation across 10 runs of the algorithm. Bottom panel: peak-to-peak difference between the measured and predicted waveforms, averaged across 10 runs of the algorithm. The solid line corresponds to this metric calculated along the estimated threshold contour; the dashed line corresponds to this metric calculated across the entire stimulus space.

The bottom panel of Fig. 9 shows the difference in peak-to-peak amplitude of the measured tone-burst ABR waveforms and the GP model predicted tone-burst ABR waveforms as a function of the initialization value of *ℓ*. As above, the data is presented as the mean and standard deviation across 10 consecutive runs of the algorithm. The solid line corresponds to this metric for the waveforms along the threshold estimate, while the dashed line corresponds to this metric calculated for all stimuli conditions. This panel demonstrates that the model is providing reasonable waveform predictions when the initialization value of *ℓ* is between [-2, 8]. This is not surprising given that the optimized value of *ℓ* is around the same value of -1 within this range and demonstrates the relative robustness of hyperparameter optimziation for a large range of initialization values. However, the initialization value that leads to the smallest standard deviation is -2, and typically increases with increasing initialization values. These simulations illustrate the robustness of the algorithm against variabilities in hyperparameter initialization. Nevertheless, issues related to over- and under-fitting could occur when the model is initialized to be too flexible (e.g., too small *ℓ* values) or too rigid (e.g., too large *ℓ* values).

#### 5.3.3. Termination criteria considerations

In the current article, the algorithm is run up to a fixed number of 20 stimuli. To investigate this termination criteria, the algorithm is run up to 40 iterations 10 times for ear M1E1. The top panel of Fig. 10 shows the average threshold estimate for each frequency as a function of iteration. The threshold estimate is relatively stable by 20 or 30 iterations, although it does continue to change slowly as more data is included in the model.

**Figure 10:**
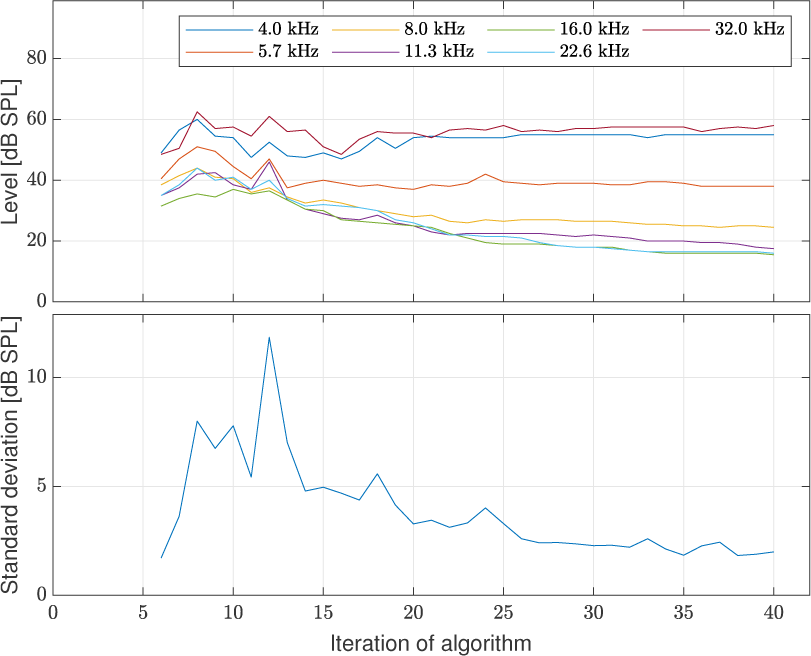
Both panels simulated for ear M1E1. Top panel: the threshold estimate for each frequency averaged across 10 runs of the algorithm as a function of iteration. Bottom panel: the standard deviation of threshold estimate across 10 runs of the algorithm, averaged over frequency.

The bottom panel of Fig. 10 shows the standard deviation across the 10 runs of the threshold estimate, averaged across frequencies. As in the top panel, the threshold estimates are relatively stable by 20 or 30 iterations, but become more similar across repeated runs of the algorithm as more iterations are included in adaptive algorithm.

## 6. Conclusion

The current article proposes an algorithm that pairs a Gaussian process model with a Bayesian adaptive stimulus selection scheme to predict ABR threshold contours across frequency. In numerical experiments using previously collected ABR waveforms, the algorithm is able to efficiently estimate ABR thresholds, leading to 3 to 5 time-savings compared to the original data collection protocols. Auditory brainstem response thresholds estimated by the adaptive algorithm match those of human raters within 10 dB for 19 out of 27 ears. Further, test/retest performance of the algorithm yields threshold estimates with less variance compared to the threshold estimates from four independent human raters.

## 7. Acknowledgments

The authors would like to thank Eileen Brister, MacKenzie Mills, Bohua Hu, Celia Zhang, Anselm Joseph Gadenstaetter, and Kenneth Henry for generously providing ABR datasets. Additionally, the authors thank Minhao Zhang and Yuxuan Wang for their contributions to the code. This work was supported by NIH grant R01DC017988 (PI: Y. Shen) and PHS GRANT NUMBER 2 T32 DC 5361-21 (Perkel D. J. and J. A. Sisneros).

